# An integrated human forebrain organoid reveals microglia-mediated CD8⁺ T cell recruitment and neuroimmune dysfunction in Alzheimer’s disease pathology

**DOI:** 10.64898/2026.05.25.727443

**Authors:** Yilin Feng, Feifei Yu, Jiyuan Tang, Honghui Zheng, Zitian Wang, Shaohua Ma

## Abstract

Genetic evidence implicates immune dysfunction in Alzheimer’s disease (AD), yet the neuroimmune mechanisms operating in human tissue remain poorly defined. Here, we establish a modular human forebrain organoid platform that systematically integrates iPSC-derived microglia and CD8^+^ T cells to reconstitute multicellular Alzheimer’s disease pathology. This system enables functional interrogation of both innate and adaptive immune components in a human-relevant context. Using this platform, we demonstrate that microglia mediate amyloid-β clearance and neuronal maturation but also drive inflammatory activation and recruit CD8^+^ T cells through CCL4/5 signaling via CCR1/5 and CXCR3, establishing a neuroinflammatory feedback loop. Pharmacological targeting of CCR5 or CXCR3 blocks T cell recruitment and modulates autophagy in a microglia-dependent manner. This modular organoid platform provides a versatile tool for dissecting neuron–immune interactions and enables cell-type-specific therapeutic screening in human neuroinflammatory disease models.

**Highlights:**

1) A modular human forebrain organoid platform integrating innate (microglia) and adaptive (CD8^+^ T cells) immunity recapitulates key features of the human neuroimmune environment
2) Enables mechanistic dissection of multicellular interactions underlying Alzheimer’s disease pathology
3) Overcomes limitations of traditional animal models by resolving human-specific, cell-type-specific neuroimmune mechanisms
4) Establishes a new approach methodology for studying neuroimmune disorders and advancing drug discovery

## Introduction

Alzheimer’s disease (AD) is featured by amyloid-beta (Aβ) accumulation, which triggers a complex cascade of glial activation, neuroinflammation, and neuronal damage^1^. While resident microglia and astrocytes drive much of this local neuroinflammatory landscape^2,3^, emerging evidence highlights the active infiltration of adaptive immune cells - particularly CD8+ T cells - into the AD brain parenchyma^4^. We previously demonstrated in transgenic mouse models that CD8+ T cells are enriched at pathological lesions and play an essential role in AD progression^5^. However, the precise mechanisms governing their recruitment and their interplay with resident central nervous system (CNS) cells remain poorly understood, largely due to profound translational gaps between murine models and human immunity.

The CCL5-CCR5 chemokine signaling axis has emerged as a critical node integrating these cellular responses^6,7^. Expressed on both CNS resident cells and peripheral immune cells, the CCR5 receptor and its glial-derived ligand CCL5 facilitate both local neuroinflammation and the chemotactic infiltration of CD8+ T cells^8^. Yet, determining the exact, cell-type-specific roles of CCL5-CCR5 and other signaling pathways^9^ in human AD has been stalled by a lack of experimental systems capable of faithfully capturing human, multicellular neuroimmune dynamics.

To bridge this fundamental gap, we engineered a modular, human forebrain organoid platform that systematically reconstructs both innate and adaptive neuroimmune microenvironments. Unlike traditional 2D cultures or in vivo murine models, this 3D system structurally and functionally integrates human induced pluripotent stem cell (iPSC)-derived forebrain organoids with iPSC-derived microglia and primary peripheral CD8+ T cells. By exposing this platform to the highly amyloidogenic Aβ42 isoform, we established an experimentally tractable, tri-cellular model that dynamically mimics the human AD neuroinflammatory milieu.

Crucially, the modular nature of this platform permits precise spatiotemporal control over cellular composition, stepwise tracking of T cell infiltration, and real-time functional readouts of neuro-glial-T cell crosstalk. Utilizing this system, we successfully recapitulated key features of human AD immune dysfunction, dissected the mechanisms translating Aβ pathology into T cell recruitment in a human cellular context, and identified and verified novel immunomodulatory cues. Ultimately, this integrated organoid platform provides a comprehensive and versatile framework for dissecting complex neuroimmune interactions and identifying cell-type-specific therapeutic targets in AD.

## Result

### Aβ42 Impairs Forebrain Organoid Maturation and Induces a Stress-Associated State

To establish a human neural substrate for modeling Alzheimer’s disease (AD), we differentiated forebrain organoids from human iPSCs, following previous established protocol^10^, and exposed them to Aβ42 oligomers for 8 days beginning at day 90 of differentiation (Fig. 1A). Immunofluorescence staining confirmed the presence of cortical progenitor and neuronal populations, including TBR2^+^, CTIP2^+^, and SOX2^+^ cells (Fig. 1B) as well as SATB2^+^, CTIP2^+^, and BRN2^+^ neurons (Fig. 1C), in both control (CTR) and Aβ42-treated organoids. Forebrain regional identity was maintained following Aβ42 exposure, as evidenced by sustained expression of FOXG1, MAP2, SOX2, and PAX6 (Fig. 1D). Astrocyte-associated populations expressing GFAP and S100B were also present in both conditions (Fig. 1E). Immunostaining with the 6E10 antibody confirmed significant Aβ accumulation in treated organoids (Fig. 1F; P < 0.0001).

**Figure 1.**
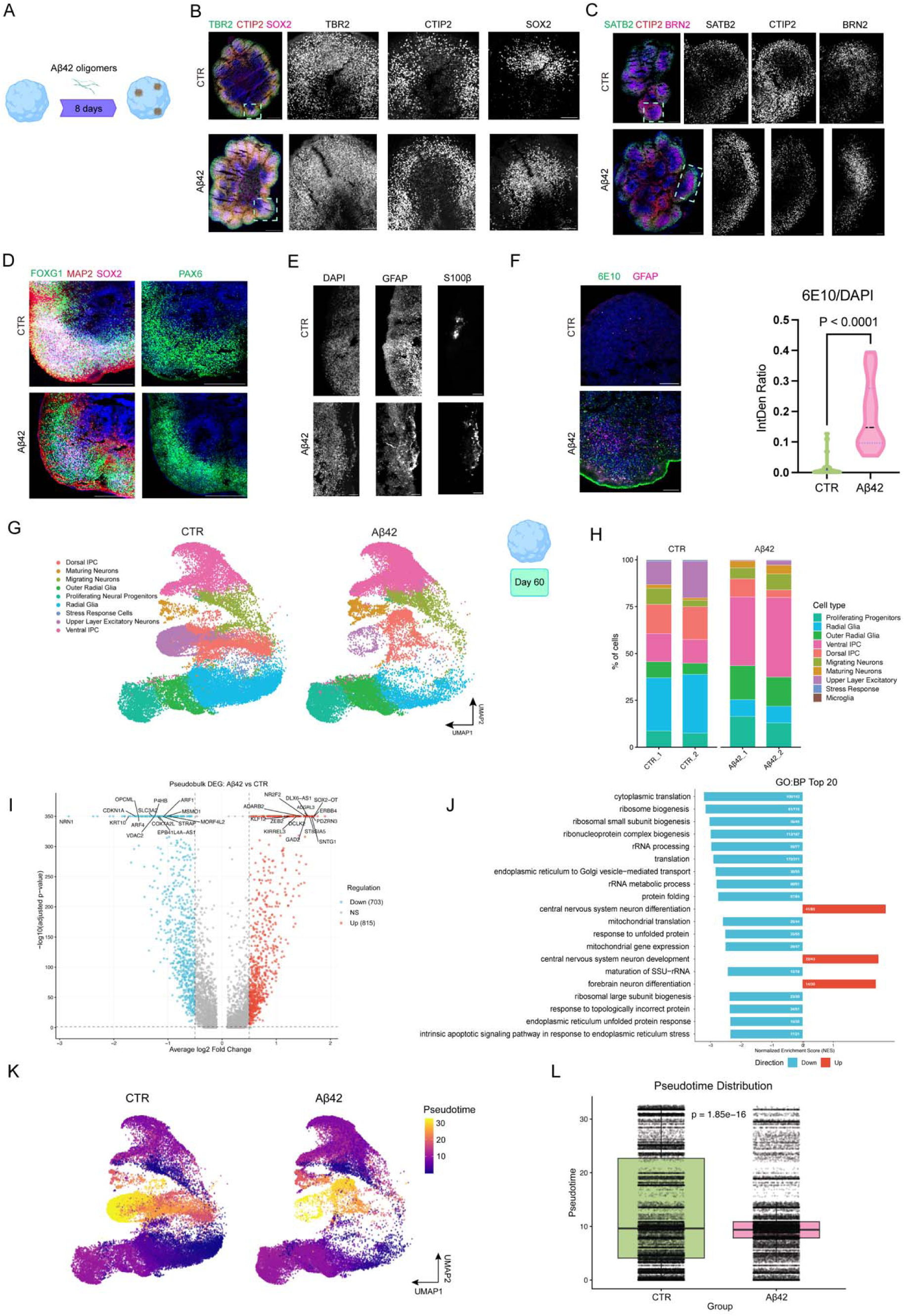
Effects of Aβ42 oligomer exposure on forebrain organoids. (A) Schematic of the experimental design. Forebrain organoids were differentiated and exposed to Aβ42 oligomers for 8 days starting at day 90. (B–C) Immunofluorescence staining of cortical progenitor and neuronal layer markers in control (CTR) and Aβ42-treated organoids: TBR2, CTIP2, and SOX2 (B); SATB2, CTIP2, and BRN2 (C). Scale bars: 500 µm (overview), 100 µm (insets). (D) Immunofluorescence staining for FOXG1, MAP2, SOX2, and PAX6 in CTR and Aβ42-treated organoids. Scale bars: 500 µm. (E) Immunostaining for GFAP and S100B in CTR and Aβ42-treated organoids. Scale bars: 50 µm. (F) Aβ detected with the 6E10 antibody in CTR and Aβ42-treated organoids. Right, quantification of 6E10 signal intensity normalized to DAPI (violin plot; unpaired t test, P < 0.0001; n = 4–6 organoids per group). Scale bars: 100 µm. (G) UMAP visualization of integrated single-cell RNA-seq data from day 60 CTR and Aβ42-treated organoids, split by condition and colored by annotated cell type. (H) Relative cell-type composition of CTR and Aβ42-treated organoids, shown as stacked bars for each sequencing library (n = 2 per condition); statistics in Supplementary Table 1. (I) Volcano plot of differentially expressed genes from pseudobulk analysis comparing Aβ42-treated organoids with CTR; selected genes are labeled. (J) Gene set enrichment analysis (GSEA) of Gene Ontology Biological Process (GO:BP) terms using differentially expressed genes from the pseudobulk comparison in (I), ordered by normalized enrichment score. (K) Pseudotime trajectories inferred with Monocle3 and projected onto UMAP embeddings for CTR (left) and Aβ42-treated (right) organoids. (L) Distribution of pseudotime values in CTR and Aβ42-treated organoids (Wilcoxon rank-sum test).

To characterize the transcriptional impact of Aβ42 exposure, we performed single-cell RNA sequencing on day 60 CTR and Aβ42-treated organoids. UMAP visualization of integrated data revealed diverse neural cell types, including radial glia, intermediate progenitor cells (IPCs), migrating neurons, maturing neurons, and upper-layer excitatory neurons (Fig. 1G), which were annotated on the basis of canonical marker expression (Fig. S1). Comparison of relative cell-type proportions showed shifts following Aβ42 treatment, including an increase in stress-responsive populations and a decrease in maturing neurons (Fig. 1H). Compositional differences were assessed by Bayesian compositional analysis (scCODA)^11^ applied to the per-library cell proportions reported in Supplementary Table 1, with maturing neurons as the reference population; credible changes are indicated in Supplementary Table 1. Pseudobulk differential expression analysis revealed upregulation of stress-associated genes and downregulation of neuronal maturation-related genes in Aβ42-treated organoids (Fig. 1I). Gene set enrichment analysis (GSEA) confirmed activation of cellular stress pathways and depletion of neuronal maturation programs (Fig. 1J). Pseudotime trajectory analysis using Monocle3 showed that Aβ42-treated organoids exhibited a significant shift toward lower pseudotime values compared with CTR (Fig. 1K–L; Wilcoxon rank-sum test), indicating that Aβ42 exposure arrests cells in an immature, stress-associated state.

Together, these data establish that Aβ42 exposure alone induces transcriptional stress and impairs neuronal maturation in human forebrain organoids, creating a vulnerable neural substrate for subsequent immune interactions.

### A 2D-3D Hybrid Organoid System Enables Functional Interrogation of A**β**42-Induced Synaptic and Network Dysfunction

To enable functional analysis of Aβ42-induced pathology, we developed a 2D-3D hybrid organoid system. Day 90 cortical organoids were either maintained as intact 3D structures for bulk RNA-seq analysis or sectioned into 300-µm slices and cultured on Matrigel-coated plates, preserving three-dimensional cytoarchitecture while providing optical access for imaging^12^ (Fig. 2B). Bulk RNA-seq of intact 3D organoids confirmed that Aβ42 treatment induced transcriptional changes consistent with our single-cell data, including upregulation of stress-associated genes and downregulation of maturation-related genes (Fig. 2A).

**Figure 2.**
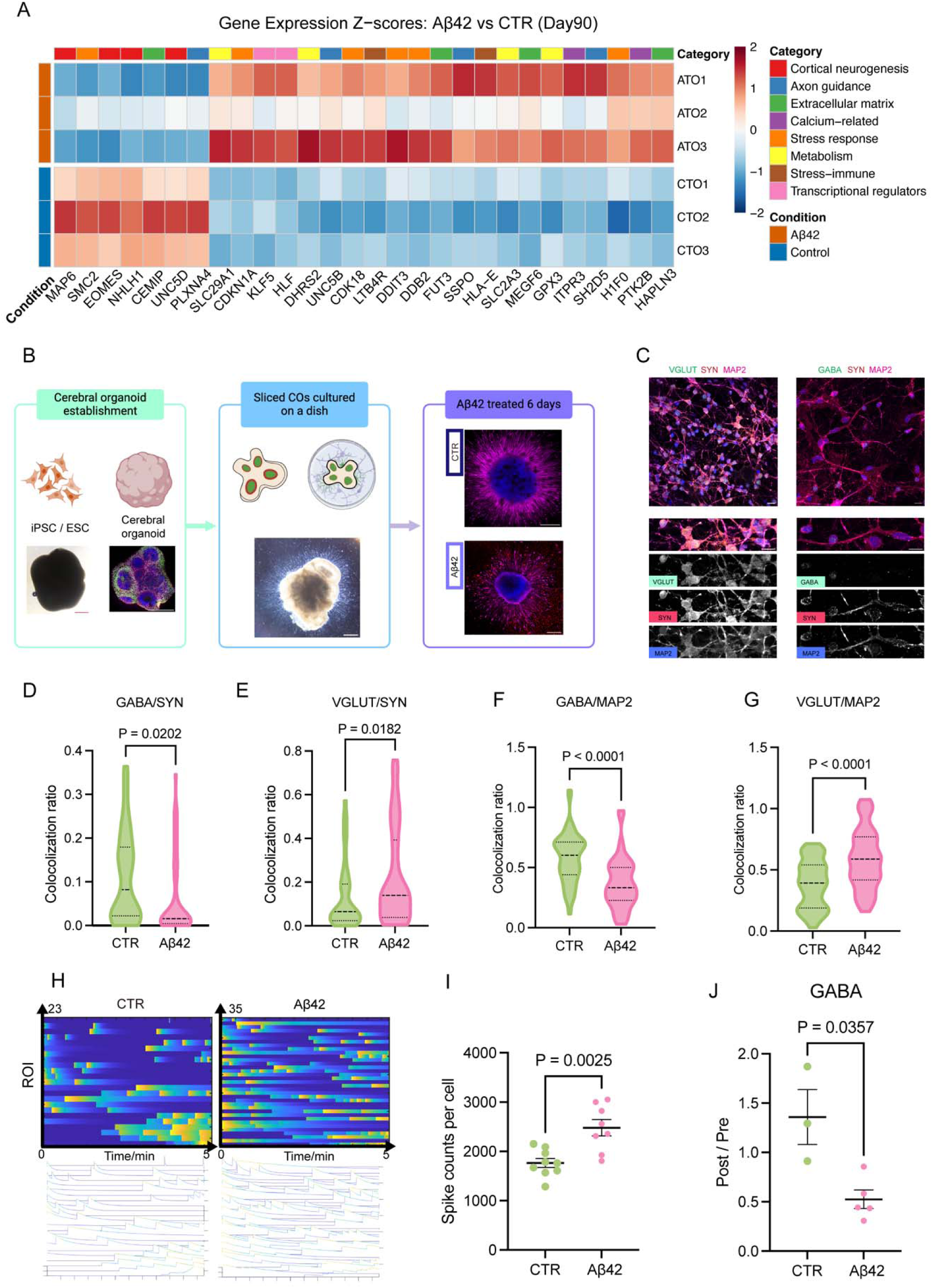
Development and characterization of a 2D–3D hybrid organoid model for analyzing Aβ42-induced pathology. (A) Heatmap of the top 30 most significantly differentially expressed genes from bulk RNA-seq of day 90 3D Aβ42-treated and CTR organoids. Expression values are shown as scaled normalized expression across individual samples, with hierarchical clustering applied to both genes and samples. (B) Experimental workflow. Day 90 cortical organoids were either maintained as intact 3D organoids for bulk RNA-seq or sectioned into 300-µm slices and cultured on Matrigel-coated plates to generate a 2D–3D hybrid organoid system for synaptic and functional analyses. Hybrid cultures were treated with 5 µM Aβ42 oligomers for 6 days prior to downstream assays. (C) Representative immunofluorescence images of synaptophysin and MAP2 co-stained with VGLUT or GABA, used to determine synapse ratios. Scale bar: 10 µm. (D–G) Quantification of synaptic marker density in CTR and Aβ42-treated 2D–3D hybrid organoids: excitatory synapses (VGLUT/synaptophysin) and inhibitory synapses (GABA/synaptophysin). (H) Representative calcium imaging traces from CTR and Aβ42-treated 2D–3D hybrid organoids, acquired at 7.67 s per frame. (I) Quantification of spike frequency per cell from calcium imaging in CTR and Aβ42-treated hybrid organoids. (J) GABA response expressed as the post-/pre-treatment spike count ratio. Data are mean ± SEM. For immunofluorescence and functional analyses, n = 4–6 organoids per group. Statistical significance was assessed using unpaired t tests.

We next used the hybrid system to assess synaptic and network-level consequences of Aβ42 exposure. Immunofluorescence staining for synaptic markers revealed a significant shift in excitatory/inhibitory balance: Aβ42-treated organoids exhibited increased density of VGLUT^+^ excitatory synapses and decreased density of GABA^+^ inhibitory synapses, normalized to synaptophysin (Fig. 2C-G). This synaptic imbalance was accompanied by functional network alterations. Calcium imaging revealed significantly increased spike frequency per cell in Aβ42-treated hybrid organoids compared with CTR (Fig. 2H-I). Furthermore, GABA response analysis showed an altered post-/pre-treatment spike count ratio in Aβ42-treated organoids (Fig. 2J), suggesting impaired inhibitory signaling.

Together, these results indicate that Aβ42 exposure shifts the excitatory–inhibitory synaptic balance and increases spontaneous network activity in human forebrain organoids, consistent with the neuronal hyperexcitability reported in early AD^13^.

### Integration of iPSC-Derived Microglia Modulates A**β**42-Associated Transcriptional States and Neural Activity

Having established a Aβ42-exposed neural substrate with defined functional deficits, we next sought to incorporate innate immunity. We generated iPSC-derived microglia (iMGLs) through sequential differentiation of hematopoietic progenitors to immature and then mature microglia (Fig. 3A). Flow cytometry confirmed that approximately 96% of cells were CD11b^+^, with ∼50% exhibiting a CD45 phenotype consistent with microglial maturation (Fig. 3B). When co-cultured with Aβ42-treated organoids, iMGLs associated with and engulfed Aβ42 oligomers, as visualized by IBA1 and 6E10 co-staining (Fig. 3B). ELISA quantification of culture media showed that iMGL integration significantly reduced Aβ42 levels compared with organoids alone (Fig. 3C), confirming functional Aβ clearance. To understand how microglia reshape the cellular landscape, we performed single-cell RNA sequencing on Aβ42-treated organoids with and without iMGL co-culture (Aβ42+iMGL and Aβ42 conditions). UMAP visualization revealed the appearance of a distinct microglial cluster in the Aβ42+iMGL group, with all major neural populations preserved (Fig. 3D). Comparison of cell-type proportions showed shifts following microglial integration, including changes in progenitor and neuronal subtypes (Fig. 3E). Pseudotime analysis showed that microglial co-culture shifted cells toward higher pseudotime values compared with Aβ42 alone (Fig. 3F–G), consistent with a shift in organoid cell-type composition toward more differentiated populations (Fig. 3E) and a partial reversal of the Aβ42-associated skew toward progenitor states. Cell-cell interaction analysis using CellChat identified extensive signaling networks between microglia and neural populations (Fig. 3H). Microglia exhibited both outgoing signals, including neurotrophic (PTN, IGF1), synaptic (NRXN3-NLGN1), and immunomodulatory (LGALS9-HAVCR2) pathways, and incoming signals from neural progenitors and neurons (Fig. 3I-J), revealing bidirectional communication.

**Figure 3.**
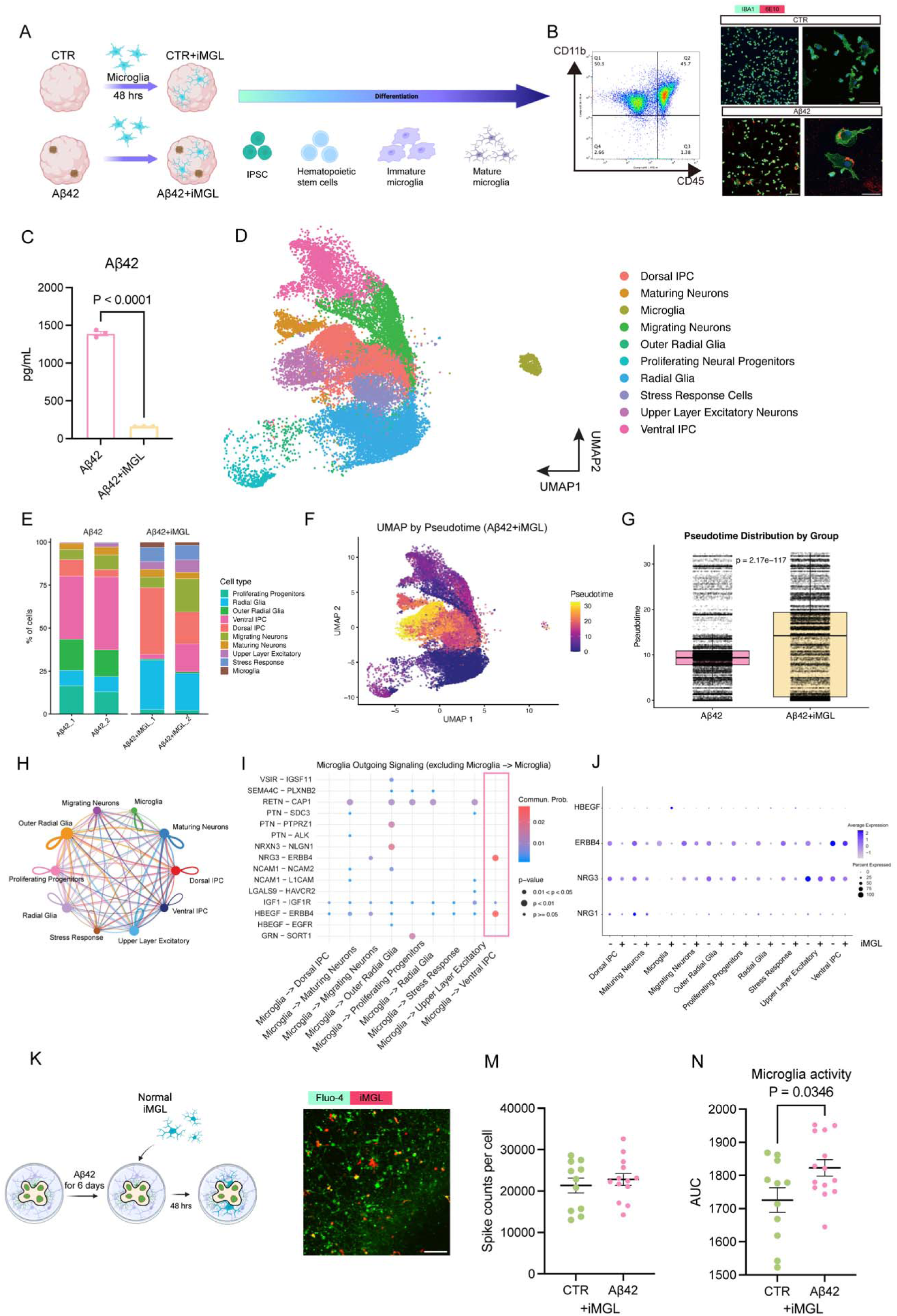
A microglia-integrated forebrain organoid model of Aβ42 exposure. (A) Experimental scheme for the microglia-integrated forebrain organoid model. Forebrain organoids were treated with vehicle (0.1% DMSO; CTR) or Aβ42 oligomers (5 µM) for 6 days, followed by 48 h of co-culture with iPSC-derived microglia (iMGLs) to generate CTR+iMGL and Aβ42+iMGL conditions. The scheme also depicts iPSC differentiation through hematopoietic progenitor, immature microglia, and mature microglia stages. (B) Characterization of iMGLs. Left, flow cytometry of CD11b and CD45 (∼96% CD11b^+^; ∼50% CD45^low). Right, representative immunofluorescence images of iMGLs stained for IBA1 (green) with Aβ42 detected by 6E10 (red). Scale bars: 100 µm (overview), 50 µm (high magnification). (C) Aβ42 concentration in culture medium measured by ELISA 48 h after iMGL integration, in Aβ42-treated organoids with and without iMGLs (unpaired t test, P < 0.0001; n = 3 samples per group). Data are mean ± SEM. (D) UMAP visualization of integrated single-cell RNA-seq data from Aβ42-treated forebrain organoids (Aβ42) and Aβ42-treated organoids co-cultured with microglia (Aβ42+iMGL), split by condition and colored by annotated cell type; the microglial cluster is shown in yellow. (E) Relative cell-type composition of Aβ42-treated organoids with and without iMGL co-culture, shown as stacked bars for each sequencing library (n = 2 per condition); statistics in Supplementary Table 1. (F) Pseudotime trajectory inferred with Monocle3 and projected onto the UMAP embedding for the Aβ42+iMGL condition. Color scale indicates pseudotime from early (purple/blue) to late (yellow/red). (G) Distribution of pseudotime values in Aβ42-treated organoids with and without iMGL co-culture (Wilcoxon rank-sum test, P = 2.17 × 10⁻¹). (H) Predicted cell–cell interaction network between cell types in Aβ42+iMGL organoids. Node size and edge thickness scale with interaction strength; colors denote cell populations. (I, J) Outgoing (I) and incoming (J) microglial signaling across pathways, relative to other cell types in the organoid. (K) Experimental design for functional analyses in the 2D–3D hybrid forebrain organoid system. Organoid slices were treated with Aβ42 (5 µM) for 6 days, followed by 48 h of co-culture with iMGLs prior to calcium imaging. (L) Representative calcium imaging field showing neuronal activity labeled with Fluo-4 (green) and iMGLs labeled with CellTracker CMTPX (red) in Aβ42-treated hybrid organoids. Scale bar: 100 µm. (M) Neuronal calcium activity (spike counts per cell) in CTR+iMGL and Aβ42+iMGL conditions (unpaired t test; n = 3–4 organoids per group). Individual dots represent single cells; horizontal lines indicate mean ± SEM. (N) Microglial calcium activity expressed as area under the curve (AUC) of ΔF/F₀ traces in CTR+iMGL and Aβ42+iMGL conditions (unpaired t test, P = 0.0346; n = 3–4 organoids per group). Individual dots represent single microglia; horizontal lines indicate mean ± SEM. 3K is created in BioRender. Feng, Y. (2026) https://BioRender.com/6o4kfkx.

We next assessed the functional impact of microglial integration using the 2D-3D hybrid system (Fig. 3K). Calcium imaging with Fluo-4-labeled neurons and CellTracker CMTPX-labeled microglia (Fig. 3L) revealed that, in the presence of microglia, the Aβ42-induced neuronal hyperactivity was normalized: there was no significant difference in spike frequency between CTR+iMGL and Aβ42+iMGL conditions (Fig. 3M). However, microglia themselves exhibited significantly increased calcium activity in the Aβ42+iMGL condition compared with CTR+iMGL (Fig. 3N), indicating that while microglia protect neurons from hyperactivity, they become activated in the Aβ environment.

These data demonstrate that microglia integrate into forebrain organoids, clear Aβ, promote neuronal maturation, and normalize network activity, yet simultaneously enter a hyperactive state, revealing their dual functionality in AD pathology.

### Microglia Drive CD8**^+^** T Cell Recruitment and Establish a Pathological Neuroimmune Tri-Cellular Unit

The persistent hyperactive state of microglia in Aβ42 environments prompted us to examine their inflammatory output. Conditioned media from CTR+iMGL and Aβ42+iMGL organoids were analyzed for cytokine and chemokine secretion. Aβ42+iMGL organoids exhibited significantly elevated levels of CCL4, CCL5, IL-1β, and TGF-β1, with no change in CXCL10 (Fig. 4A-E). These chemokines are known chemoattractants for adaptive immune cells, particularly CD8^+^ T cells.

**Figure 4.**
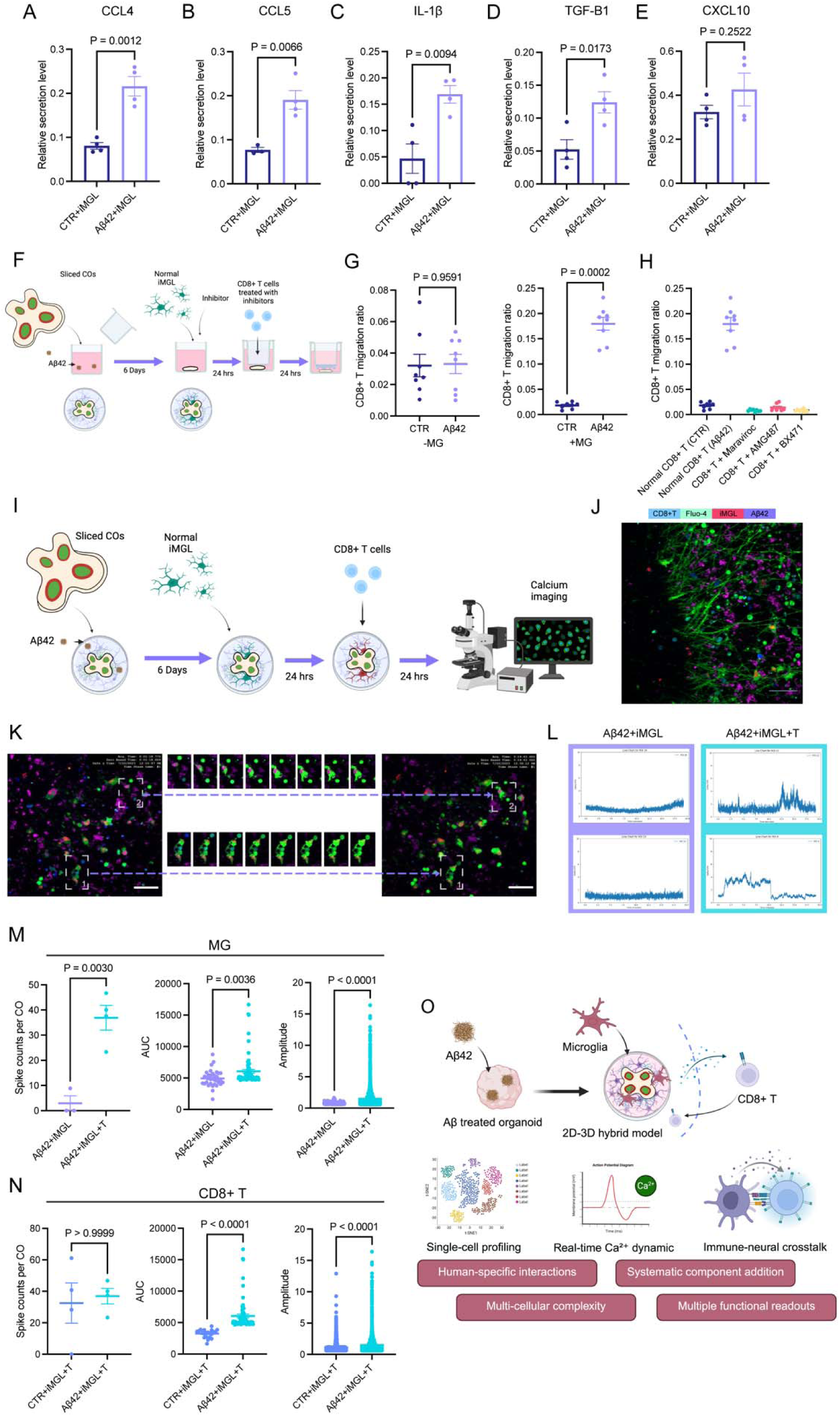
Cytokine secretion, CD8^+^ T cell migration, and microglia–T cell calcium activity in microglia-integrated Aβ42-treated forebrain organoids. (A–E) Relative secretion of CCL4 (A), CCL5 (B), IL-1β (C), TGF-β1 (D), and CXCL10 (E) measured in conditioned media from CTR+iMGL and Aβ42+iMGL forebrain organoids. Data are mean ± SEM; unpaired t tests. (F) Experimental schematic of the 2D–3D hybrid forebrain organoid model used to assess CD8^+^ T cell migration. Organoids were treated with Aβ42 for 6 days and co-cultured with iPSC-derived microglia (iMGLs), followed by transwell migration assays using CD8^+^ T cells with or without chemokine receptor inhibitors targeting CCR5, CXCR3, and CCR1. (G) Quantification of CD8^+^ T cell migration toward CTR and Aβ42-treated organoids in the absence (−MG) or presence (+MG) of iMGL co-culture. (H) CD8^+^ T cell migration toward Aβ42+iMGL organoids following blockade of CCR5, CXCR3, or CCR1 on CD8^+^ T cells, compared with the untreated Aβ42+iMGL condition. (I) Experimental design for calcium imaging of microglia–T cell interactions. iMGLs were integrated into Aβ42-treated organoids on day 6, CD8^+^ T cells were added on day 7, and calcium imaging was performed on day 8. (J) Representative multi-channel image showing CD8^+^ T cells (CellTrace Violet, blue), intracellular calcium signal (Fluo-4 AM, green), iMGLs (CellTracker CMTPX, magenta), and Aβ42 (647 nm). Scale bar: 50 µm. (K) Time-lapse calcium imaging of iMGLs in the presence of CD8^+^ T cells over a 13-min window. Insets show individual iMGLs in direct contact with T cells or associated with Aβ42. Scale bar: 50 µm. (L) Microglial calcium activity in Aβ42+iMGL and Aβ42+iMGL+T conditions, quantified as spike count per cell, area under the curve (AUC), and spike amplitude. Individual dots represent single iMGLs. (M) Microglial calcium activity in CTR+iMGL+T and Aβ42+iMGL+T conditions, using the same metrics as in (L). (N) CD8^+^ T cell calcium activity in CTR+iMGL+T and Aβ42+iMGL+T conditions, quantified as spike count per cell, AUC, and spike amplitude. Calcium imaging data were acquired at 3.8-s intervals. (O) Schematic of the integrated human forebrain organoid platform, in which Aβ42-treated organoids are converted to a 2D–3D hybrid system permitting stepwise integration of iPSC-derived microglia and CD8^+^ T cells, with readouts comprising single-cell transcriptional profiling, real-time calcium imaging, and assays of immune–neural cellular interaction. Data are mean ± SEM; statistical significance was assessed using unpaired t tests. For cytokine analyses, n = 3 samples per group. For migration and calcium imaging analyses, n = 3–4 organoids per group, with individual dots representing single cells where indicated. 4O is created in BioRender. Feng, Y. (2026) https://BioRender.com/wgpmokj.

To test whether microglial-derived signals recruit CD8^+^ T cells, we established a transwell migration assay using the 2D-3D hybrid system (Fig. 4F). In the absence of microglia, CD8^+^ T cells showed no significant migration toward either CTR or Aβ42-treated organoids (Fig. 4G). However, when microglia were present, Aβ42-treated organoids induced significantly increased T cell migration compared with CTR+iMGL (Fig. 4G). This recruitment was mediated by chemokine receptors on T cells: pharmacological inhibition of CCR5, CXCR3, or CCR1 significantly reduced migration toward Aβ42+iMGL organoids (Fig. 4H).

We next established a tri-culture system to visualize and quantify microglia-T cell interactions (Fig. 4I). Forebrain organoids were treated with Aβ42 for 6 days, co-cultured with iMGLs for 48 hours, and then CD8^+^ T cells were added 24 hours prior to calcium imaging. Multi-channel imaging enabled simultaneous visualization of T cells (CellTrace Violet), calcium activity (Fluo-4 AM), microglia (CellTracker CMTPX), and Aβ42 (647 nm) (Fig. 4J). Time-lapse imaging revealed heterogeneous microglial activation states, with some microglia engaging in direct T cell interactions and others exhibiting Aβ42-associated activity (Fig. 4K).

Quantitative analysis showed that the presence of T cells further amplified microglial hyperactivity: Aβ42+iMGL+T organoids exhibited significantly increased microglial spike count, area under the curve (AUC), and spike amplitude compared with Aβ42+iMGL without T cells (Figs. 4L, 4M). Microglial activity was also significantly higher in Aβ42+iMGL+T compared with CTR+iMGL+T (Fig. 4N), confirming that the Aβ environment drives sustained microglial activation even in the presence of T cells. CD8^+^ T cells themselves exhibited calcium activity that was modulated by the environment, with trends toward increased activity in the Aβ42 condition (Fig. 4N).

Together, these data establish a tri-cellular neuroimmune unit in which: (1) Aβ42 creates a stressed neural substrate, (2) microglia respond by clearing Aβ and promoting maturation but become hyperactive and secrete inflammatory chemokines, (3) these chemokines recruit CD8^+^ T cells via CCR5/CXCR3 signaling, and (4) recruited T cells further amplify microglial activation, creating a positive feedback loop of neuroinflammation (summarized in Fig. 4O).

This modular human forebrain organoid platform, incorporating both innate and adaptive immunity, enables systematic dissection of multicellular neuroimmune interactions in AD and provides an experimentally tractable system for identifying cell-type-specific therapeutic targets.

## Discussion

The neuroimmune axis is central to AD pathogenesis, yet the human cellular mechanisms translating amyloid pathology into sustained neuroinflammation remain poorly understood. By establishing a modular human forebrain organoid platform that sequentially integrates innate and adaptive immunity, we successfully deconstructed the tri-cellular interactions underlying AD neuroinflammation. This stepwise approach revealed a multi-stage positive feedback loop: (i) Aβ42 exposure induces neuronal stress; (ii) microglia clear Aβ (Fig. 3B-C) and normalize neuronal hyperactivity (Fig. 3M) but enter a chronically active, chemokine-secreting state (Fig. 3N, 4A-E); (iii) these microglial signals recruit CD8^+^ T cells via CCR5/CXCR3 pathways (Fig. 4H); and (iv) the recruited T cells subsequently amplify microglial hyperactivity (Fig. 4L). Crucially, T cells failed to migrate to Aβ-treated organoids in the absence of microglia (Fig. 4G). This provides a mechanistic demonstration in human-derived cells that microglial activation serves as the essential bridge between amyloid pathology and the adaptive immune recruitment recently documented in human AD brains.

Furthermore, our findings experimentally resolve the functional duality of microglia in AD. Within our platform, microglia simultaneously executed protective functions—engulfing Aβ and promoting neuronal maturation (Fig. 3F-G)—while initiating pathological inflammatory cascades. This duality indicates that microglial activation is not a binary state: within the Aβ42 environment, clearance and neuroprotective functions are detectable alongside inflammatory, T-cell-recruiting outputs at the same timepoint. We identified the CCL5-CCR5 axis as the master integrator of this neuroimmune transition. Microglia-derived CCL5 directly drives CD8^+^ T cell chemotaxis, while cell-cell interaction analyses revealed extensive CCR5-associated signaling with neurons and progenitors (Fig. 3H-J). Inhibiting CCR5 successfully blocked T cell recruitment (Fig. 4H) but also impacted cell-specific autophagy, indicating that CCR5 acts as a pleiotropic regulator of neuroimmune function rather than a simple chemoattractant. Repurposing clinical CCR5 antagonists (e.g., Maraviroc) or CXCR3 inhibitors offers an immediate therapeutic avenue to interrupt this pathological cascade^14,15^; however, our data caution that precise modulation is necessary to suppress inflammatory outputs without sacrificing the protective clearance and maturation functions of microglia.

While our 2D-3D hybrid platform captures human-specific cellular features, transcriptomic signatures, and real-time multicellular interactions unmatched by rodent or conventional in vitro models, we acknowledge several limitations. The current system lacks vasculature and additional peripheral immune populations, utilizes a single iPSC line, and reflects an acute timeline rather than decades-long disease progression. Future iterations incorporating multi-donor genetic variability - particularly AD risk variants - and extended culture periods will be critical to fully modeling disease evolution.

The modular, tri-cellular platform described here has implications beyond AD. The same approach could be applied to model other neurodegenerative and neuroinflammatory conditions, including Parkinson’s disease, amyotrophic lateral sclerosis, multiple sclerosis, and neuropsychiatric disorders with immune components. Microglia-containing midbrain organoids have already been used to examine glial contributions to dopaminergic neuron function^16^, iPSC-based microglia–motor neuron co-cultures have been applied to ALS ^17^ and myelinating oligodendrocytes have been generated in human cortical spheroids, providing a substrate for modeling demyelinating disease^18^. Transplantation-based neuroimmune organoid models further demonstrate that microglial phenotypes are shaped by their tissue environment^19^. By enabling systematic addition of specific immune populations — for example, different T cell subsets, B cells, or peripheral monocytes — this platform can be adapted to address diverse questions about neuroimmune interactions in health and disease.

From a therapeutic perspective, our findings identify multiple intervention points. CCR5 and CXCR3 antagonists, already in clinical use or development for other indications, could be tested for their ability to block T cell infiltration and interrupt the neuroinflammatory feedback loop. Strategies to preserve microglial protective functions (Aβ clearance, neuronal maturation) while suppressing their pathological outputs (chemokine secretion, T cell recruitment) represent a particularly attractive therapeutic goal. The platform’s compatibility with high-content imaging and multi-well formats positions it for medium-throughput drug screening, and its human relevance may improve translation compared with traditional animal models.

Despite these constraints, the identification of a tri-cellular positive feedback loop offers a mechanistic explanation for the persistence of neuroinflammation in AD, even after initial pathological triggers may have subsided. Beyond AD, this modular, high-content-compatible platform provides a versatile framework to systematically dissect neuroimmune complexity, enabling the stepwise addition of distinct immune subsets to accelerate targeted drug discovery across a broad spectrum of neurodegenerative and neuropsychiatric disorders.

## Methods

### Cerebral organoids, cell lines, medium and reagent

The protocol for the establishment of cerebral organoids was adapted from methods described in our previous work^20^. All experiments in this study used the iPSC line DYR0100, acquired from the National Collection of Authenticated Cell Cultures (Catalog # SCSP-1301). Forebrain organoids and iPSC-derived microglia (iMGLs) were both generated from DYR0100, such that all organoid–microglia co-cultures are isogenic. CD8^+^ T cells were isolated from primary human PBMCs (see below) and are therefore donor-derived. A material transfer agreement (MTA) for the DYR0100 line has been duly executed. The media and reagents employed for culturing this line and for cerebral organoid development are detailed in our prior work.

### iPSC-derived microglia

Both induced and embryonic microglia were generated following a two-step induction process. First, hematopoietic cells were derived using the STEMdiff™ Hematopoietic Kit (STEMCELL Technologies, Catalog # 05310_C). These cells were then differentiated into microglia using the STEMdiff™ Microglia Differentiation Kit (Catalog # 100-0019) and matured using the STEMdiff™ Microglia Maturation Kit (Catalog # 100-0020_C). All procedures were performed in strict accordance with the manufacturer’s protocols to ensure proper induction and maturation of microglia.

### 2D/3D neural-immune model establishment in A**β**42 context

Cerebral organoids aged 90 days were sliced into 300 μm sections using a vibratome, following the methodology described in our previous work. The sliced COs were then cultured on 1% Matrigel-coated 12-well confocal plates, with one slice per well, to facilitate immunofluorescence imaging. Within 24 hours of culturing, neurons and astrocytes migrated outwards from the center of the COs. The culture was supplemented with 5 μM β-amyloid (1-42) (MedChemExpress, Catalog # HY-P1363), and the medium was changed every two days. After 5 days, 1×105 induced microglia (iMGLs) were added to each well to establish the neural immune model. The samples were fixed for further analysis 48 hours after the introduction of iMGLs.

### Isolation of human PBMC and CD8+ T cells

Blood specimens were collected from healthy donors at Zhangzhou People’s Hospital. Peripheral blood was collected from individuals over 18 years of age in 10-ml tubes containing 5 mM EDTA. The samples were transported on ice and processed within 1-3 hours of collection. PBMCs were isolated using density gradient centrifugation with Lymphoprep™ density gradient medium (1.077 g/mL). The PBMCs were then handled with care to maintain their viability and integrity.

The human CD8+ T cells were isolated from the PBMCs using the EasySep™ Direct Human CD8+ T Cell Isolation Kit (STEMCELL Technologies, Catalog # 19663) following the manufacturer’s protocols to ensure efficient and consistent isolation of CD8+ T cells.

### Flow cytometry

Validation of isolated and migrated CD8+ T cells was conducted using APC/Cyanine7 anti-human CD8 Antibody (Biolegend, Catalog # 344714). For the verification of mature microglia, FITC anti-human CD45 Antibody (Biolegend, Catalog # 304006) and PE anti-human CD11b Antibody (Biolegend, Catalog # 301306) were utilized. The compensation matrix was established and positive peak gates were set based on the respective isotype controls. The isotype controls used were APC/Cyanine7 Mouse IgG1, κ Isotype Ctrl Antibody, Catalog # 400127; PE Mouse IgG1, κ Isotype Ctrl Antibody, Catalog # 400111; FITC Mouse IgG1, κ Isotype Ctrl Antibody, Catalog # 400107.

The cells were incubated with the specified antibodies for 40 minutes on ice to ensure adequate labeling. Before analysis, the cells underwent filtration through a 300-mesh Falcon centrifuge tube to eliminate aggregates and ensure a single-cell suspension. All flow cytometry data were then analyzed using FlowJo™ Software v10.10. for detailed phenotypic characterization.

### Cell Tracker staining

Before conducting calcium imaging, Cell Tracker dyes were used to stain CD8+ T cells and iMGLs for better identification. CD8+ T cells were stained with CellTrace™ Violet (dilution 1:1000, Invitrogen, Catalog # C34557), and iMGLs were labeled with CellTracker™ CMTPX (dilution 1:1000, Invitrogen, Catalog # C34552). The cells were then incubated in the staining solution at 37 °C for 30 minutes. After incubation, the staining solution was removed, and the cells were cultured for future use within the next week.

### Human inflammatory cytokine array

For each experimental group, 8 ml of medium was collected and concentrated using Ultrafiltration Centrifugal Tubes (Millipore, Catalog # UFC903096). The medium was reduced to a volume of 200 μl through ultrafiltration. This concentrated sample was then used to perform the Cytokine Array analysis using the Human Cytokine Antibody Array (Membrane, 42 Targets) kit (Abcam, Catalog # ab133997), strictly following the manufacturer’s protocols.

Raw images of the cytokine arrays have been made available in the source data, which can be accessed via the provided Github link: https://github.com/FengYilinElaine/2-3D-AD-oranoids.

### Immunohistochemistry

Validation of COs was based on previous work. The primary antibodies used in this study are listed in Supplementary Table 2.

### Quantitative analysis of A**β**42 levels

Quantitative analysis of Aβ42 levels was performed using the Amyloid beta 42 Human ELISA Kit (Invitrogen, Catalog # KHB3441).

### Bulk RNA sequencing analysis

Bulk RNA sequencing analysis was performed on 3D cerebral organoids, iMGLs and eMGLs, as described in the results section. The samples were collected quickly and flash-frozen in liquid nitrogen to preserve the RNA integrity. After freezing, the samples were transported on dry ice to BGI for RNA extraction and sequencing. All RNA analyses, including data processing and interpretation, were conducted on the Dr. Tom platform (BGI).

### Single-cell RNA-seq Library Preparation and Sequencing

For isolation of iPSC-derived forebrain cells, cells were collected from CTR and Aβ42-treated organoids at day 60, and from Aβ42-treated organoids with and without a 48-hour iMGL co-culture (Aβ42 and Aβ42+iMGL). Two biological replicates were included in each group. Three organoids were pooled within each replicate and washed three times with pre-chilled PBS. Single-cell suspensions were prepared by incubating with 0.5 mg/mL papain for 30 min at 37 °C in a water bath. Suspensions were filtered through a 70 µm cell strainer (BD Falcon, 352350). After centrifugation at 500 g for 5 min, the cell pellet was resuspended in PBS + 0.01% BSA.

The MGI DNBelab C-TaiM4 platform utilizes microfluidic technology to encapsulate single cells with mRNA capture magnetic beads containing unique cell barcodes (Cell Barcodes) and unique molecular identifiers (UMIs) within oil droplets. The platform employs a dual-magnetic bead system (mRNA capture beads and droplet-identification beads) to enhance capture efficiency and accuracy. Then, cell lysis was completed in the droplet, and mRNA molecules were captured by Oligo (dT) on the magnetic beads and reverse transcribed to generate cDNA with cell barcodes and UMIs.

For library construction, the cDNA was purified, amplified, and ligated with sequencing adapters to generate the final sequencing library. Library quality was checked using an Agilent 2100 Bioanalyzer (Agilent Technologies, USA). Finally, qualified libraries were sequenced in PE100 mode using DNBSEQ-T7 from BGI. The cDNA library sequencing depth was approximately 450M Reads/library, and the Oligo library sequencing depth was approximately 50M Reads/library. scRNA-seq were performed by Hangzhou Astrocyte Technology Co,.Ltd.

### scRNA-seq Data Pre-processing and Quality Control

Using the official BGI software DNBC4Tools with parameter settings, we performed filtering, alignment, quantification, and cell identification/recovery of raw data. After transcriptomic normalization processing, gene expression matrix files for each cell were obtained. The resulting gene expression matrices were imported into R (v.4.4.1) and processed using the Seurat package (v.5.3.1) for integration and quality control. rigorous filtering criteria were applied to retain high-quality cells: (1) 200 < total detected genes < 12,500; (2) total unique molecular identifier (UMI) counts > 500; (3) log10(GenesPerUMI) > 0.8; and (4) mitochondrial mRNA content < 20% and ribosomal mRNA content < 30%.

### Data analysis: Colocalization ratio of images

Confocal images were initially processed by selecting regions of interest (ROIs) and applying adaptive thresholding for binarization. Different adaptive threshold methods were used for various antibodies. The binarized images were then batch-processed using the ’Colocalization Finder’ plugin in Fiji. A script for processing colocalization in Fiji, along with detailed methods, is available at the provided GitHub link.

### Data analysis: Excitatory synapse and inhibitory synapse

The images underwent preliminary processing, which involved selecting regions of interest (ROIs) and applying adaptive thresholding techniques. Synapse numbers were quantified using the ’Puncta Analyzer’ plugin in Fiji.

### Calcium imaging and analysis

Brain organoids were cultured using the microwell air-liquid interface attachment method 43. Perform two medium changes using BrainPhys™ Neuronal Medium before beginning incubation with Flou-4 calcium indicator (Invitrogen, Catalog # F14201). Using the complete culture medium, prepare a 5 µM Flou-4 solution, add 200 µL to a single well and incubate for 2 hours in the incubator. Make two medium changes and add 500 μL of BrainPhys™ Neuronal Medium (STEMCELL Technologies, Catalog # 05790) 30 minutes prior to baseline imaging. Baseline imaging was performed using a confocal microscope (NikonA1R) for 5 minutes with no delay between frames, and the frame rate was 7.67.

After baseline imaging, neurotransmitter drugs, GABA (Sigma-Aldrich, Catalog # 56-12-2) 100 μM, were added for final concentration. After 1 minute incubation, change the medium twice and add 200 μL BrainPhys™ Neuronal Medium. Immediately image, keeping the field of view and parameters the same as the baseline image.

For batch processing in calcium imaging data analysis, CNMF-E44 was used. The code for parameters (getmat.m) can be found in the source. The sum of neuron.S, representing all neurons in the view, is recorded as the total number of spikes after processing. The number of active cells is recorded as the number of rows of neuron.S, i.e. the ROI where the spike is detected. The calcium trace overlay plot is presented by CNMFeClustering^21^. Calcium imaging analysis of microglia was initially conducted by isolating each microglial cell based on the maximum intensity projection of image sequences in the Cell Tracker channel. Subsequently, the Fluo-4 AM sequence image data for each isolated microglia was analyzed. Spikes were defined and quantified using a specific code, which is provided for reference.

### Statistical analysis

All statistical analyses in this study were conducted using commercially available software, namely Prism and Excel. Data from replicates are presented as mean ± standard error of the mean (SEM). The methodology for analyzing differences between multiple groups is detailed in the figure legends accompanying each graph. For the creation of graphical representations, Adobe Illustrator 2024 and Biorender were utilized.

### Data availability

Single-cell RNA-seq data have been deposited at GEO under accession GSE339602 and bulk RNA-seq data under accession GSE339404; both are publicly accessible as of the date of publication. Microscopy data are available at Zenodo (DOI: 10.5281/zenodo.21686712). All other raw data, including cytokine array images, and all original code are available at https://github.com/FengYilinElaine/2-3D-AD-oranoids.

### Code availability

All original code has been deposited at Zenodo and Github and is publicly available as of the date of publication.

## Acknowledgments

The work was supported by the National Key Research and Development Program of China (2024YFA0919800); Shenzhen Medical Academy of Research and Translation, SMART, B2402009; National natural science foundation of China (32371470); Key-Area Research and Development Program of Guangdong (2023B0909020003), and Shenzhen Natural Science Foundation (QNXMA20250701093300001, JCYJ20241202123909013).

## Author contributions

S.M. conceived and supervised the work. Y.F. wrote the manuscript and generated the figures. Y.F., F.Y., H.Z. and J.T. performed the experiments. Y.F., H.Z., J.T., P.E.L. and S.M. edited the final manuscript.

## Declaration of interests

The authors declare no competing interests.

## Notes

### Competing Interest Statement

The authors have declared no competing interest.

### Summary of Updates

The revised manuscript addresses reviewer concerns across several key areas: 1. Abstract & Language Moderation. CXCL10 was removed from the chemokine signaling description (Fig. 4E showed no significant change), leaving “CCL4/5 signaling via CCR1/5 and CXCR3.” “Human-specific” was replaced with “human-derived” or “in a human cellular context” throughout the Abstract, Introduction, and Discussion. The “first human-specific mechanistic demonstration” claim was removed. 2. Day 90→60 Correction. The scRNA-seq timepoint was corrected from "day 90" to "day 60" in the Results text to match Figure 1G. 3. Compositional Statistics (scCODA). Bayesian compositional analysis (scCODA) was added for cell-type proportion comparisons (Figs. 1H, 3E), with results in Supplementary Table 1. P-value notation was removed; individual library compositions are now shown as separate stacked bars. 4. Cell-Type Annotation Validation. Supplementary Figure S1 (canonical marker expression violin plots) was added to support cluster annotations, referenced in the Results. 5. Microglia Maturation Claim Moderated. The claim that "microglia promote neuronal maturation and partially reverse the Aβ42-induced immature state" was revised to describe a shift in cell-type composition toward more differentiated populations, acknowledging that trajectory inference alone cannot support the maturation claim. 6. Methods Rewrites. The scRNA-seq library preparation was completely rewritten, by removing erroneous references to cholesterol/semaglutide-treated day-30 organoids and correctly describing CTR and Aβ42-treated organoids at day 60 with/without iMGL co-culture, 3 organoids pooled per replicate. The iPSC line section was clarified: all experiments used DYR0100 only (H9 and DXR0109B removed), with isogenic organoid-microglia co-cultures explicitly stated. The Animals section was removed. 7. Data Availability. GEO accession numbers (GSE339602 for scRNA-seq, GSE339404 for bulk RNA-seq) and Zenodo DOI (10.5281/zenodo.21686712) were added. Microscopy data are now publicly deposited rather than “available upon request.” 8. Reference List Overhaul. All placeholder citations (“(ref)”, “(cell reports 2023)”) were replaced with properly formatted references. The reference list was renumbered and corrected. 9. Figure Legends. Figure titles were revised (e.g., Fig. 1 → “Effects of Aβ42 oligomer exposure on forebrain organoids”; Fig. 3 → “A microglia-integrated forebrain organoid model of Aβ42 exposure”). Interpretive statements were moved from legends to Results text. “Extended Table 1” was renamed “Supplementary Table 2.”

