## Supplemental Figures for "An integrated human forebrain organoid reveals microglia-mediated CD8⁺ T cell recruitment and neuroimmune dysfunction in Alzheimer’s disease pathology"

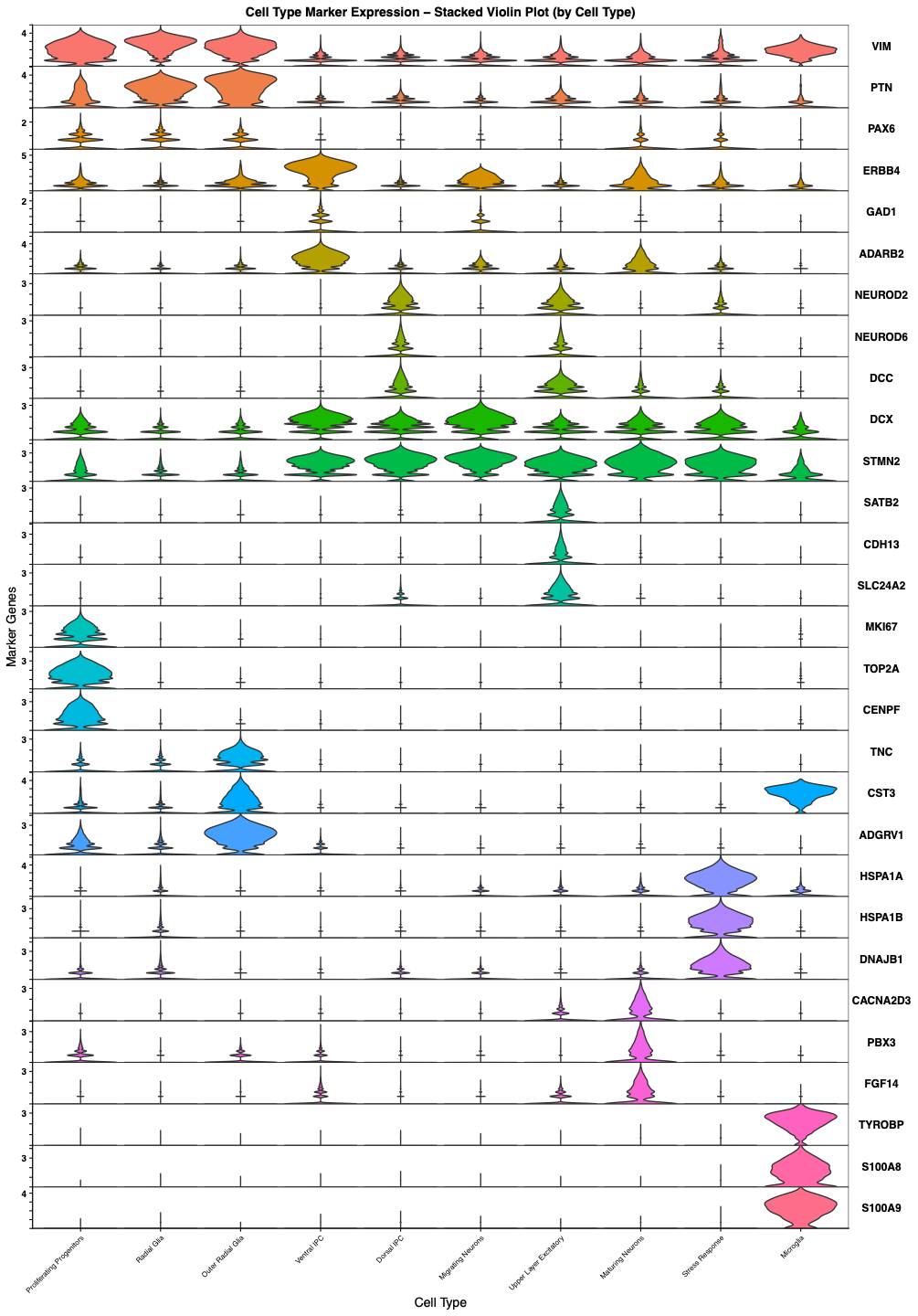

Figure S1. Cell-type annotation of forebrain organoid scRNA-seq data based on canonical marker expression.

Supplementary Table 1. scCODA compositional analysis of cell-type proportions in forebrain organoid scRNA-seq data.

Aβ42 vs CTR

| Covariate | Cell Type | Final Parameter | HDI 3% | HDI 97% | SD | Inclusion probability | Expected Sample | log2-fold change |
| --- | --- | --- | --- | --- | --- | --- | --- | --- |
| C(group, Treatment('CTR'))[T.AD] | Dorsal IPC | -1.4919004 | -2.253 | -0.736 | 0.447 | 0.98186667 | 1206.94962 | -1.1056355 |
| C(group, Treatment('CTR'))[T.AD] | Radial Glia | -1.7624079 | -2.481 | -1.076 | 0.376 | 0.99933333 | 1674.62422 | -1.4958952 |
| C(group, Treatment('CTR'))[T.AD] | Upper Layer Excitatory | -2.7579328 | -3.896 | -1.698 | 0.591 | 1 | 319.519764 | -2.9321341 |

Aβ42+iMGL vs Aβ42

| Covariate | Cell Type | Final Parameter | HDI 3% | HDI 97% | SD | Inclusion probability | Expected Sample | log2-fold change |
| --- | --- | --- | --- | --- | --- | --- | --- | --- |
| C(group, Treatment('AD'))[T.AD_mgL] | Outer Radial Glia | -2.8829225 | -4.234 | -1.52 | 0.756 | 0.9968 | 320.109792 | -2.8003052 |
| C(group, Treatment('AD'))[T.AD_mgL] | Proliferating Progenitors | -2.2364547 | -3.627 | -1.042 | 0.745 | 0.9804 | 529.605236 | -1.8676493 |
| C(group, Treatment('AD'))[T.AD_mgL] | Stress Response | 1.65606486 | -0.073 | 3.089 | 0.969 | 0.848 | 1370.11698 | 3.74806934 |
| C(group, Treatment('AD'))[T.AD_mgL] | Ventral IPC | -2.4594642 | -3.489 | -1.445 | 0.571 | 0.99486667 | 1181.00654 | -2.189384 |

Supplementary Table 1. Primary antibody list

| Name | Species | Dilution | Vendor | Catlog # |
| --- | --- | --- | --- | --- |
| SOX2 | Goat | 1:500 | R&D | AF2018 |
| CTIP2 | Rat | 1:100 | Abcam | ab18465 |
| BRN2 | Mouse | 1:300 | Millipore | MABD51 |
| SATB2 | Rabbit | 1:100 | Abcam | ab34735 |
| GFAP | Chicken | 1:1000 | Abcam | ab4674 |
| LAMININ | Rabbit | 1:1000 | Abcam | Ab30320 |
| pVIMENTIN | Mouse | 1:500 | MBL | D076-3 |
| CUX1 | Mouse | 1:100 | Santa Cruz Biotech | sc-514008 |
| NeuN | Mouse | 1:100 | Abcam | ab104224 |
| FOXG1 | Rabbit | 1:1000 | Abcam | ab18259 |
| VGLUT1 | Mouse | 1:1000 | Millipore | MAB5502 |
| MAP2 | Chicken | 1:1000 | Abcam | ab5392 |
| 6E10 | Mouse | 1:1000 | Biolegend | SIG-39300 |
| Iba1 | Rabbit | 1:200 | Wako | 019-19741 |
